# AMPylation is a specific lysosomal protein posttranslational modification in neuronal maturation

**DOI:** 10.1101/2021.03.02.433531

**Authors:** Tobias Becker, Cedric Cappel, Francesco Di Matteo, Giovanna Sonsalla, Ewelina Kaminska, Fabio Spada, Silvia Cappello, Markus Damme, Pavel Kielkowski

**Affiliations:** LMU Munich, Department of Chemistry, Butenandtstr. 5-13, 81377, Munich, Germany; University of Kiel, Institute of Biochemistry, Olshausenstr. 40, 24098, Kiel, Germany; Max Planck Institute of Psychiatry, Kraepelinstraße 2., 80804, Munich, Germany; International Max Planck Research School for Translational Psychiatry (IMPRS-TP); LMU Munich, Department of Physiological Genomics, Biomedical Center (BMC), Großhadernerstr. 9, 82152, Planegg, Germany; Helmholtz Zentrum München, Institute for Stem Cell Research, Ingolstädter Landstr. 1, 85764, Neuherberg; Graduate School of Systemic Neurosciences (GSN)

## Abstract

Protein AMPylation is a pervasive posttranslational modification with an emerging role in neurodevelopment. In metazoans the two highly conserved protein AMP-transferases together with a diverse group of AMPylated proteins have been identified using chemical proteomics and biochemical techniques. However, the function of this modification remains largely unknown. Particularly problematic is the localization of thus far identified AMPylated proteins and putative AMP-transferases. Here, we uncover protein AMPylation as a novel posttranslational modification of luminal lysosomal proteins characteristic in differentiating neurons. Through a combination of chemical proteomics, advanced gel-based separation of modified and unmodified proteins and activity assay, we show that an AMPylated, lysosomal soluble form of exonuclease PLD3 increases dramatically during neuronal maturation and that AMPylation inhibits its catalytic activity. Together, our findings unveil so far unknown lysosomal posttranslational modification, its connection to neuronal differentiation and putatively provide a novel molecular rationale to design of therapeutics for lysosomal storage diseases.

## Introduction

Protein posttranslational modifications (PTMs) provide the cell with mechanism to swiftly react on internal and external clues to maintain the cellular homeostasis. Decline of the protein homeostasis (proteostasis) is a hallmark of many neurodegenerative disorders (Hipp 2019). Regulation of protein function by PTMs include modulation of protein’s catalytic activity, localization or protein-protein interactions (Aebersold et al., 2018). Protein AMPylation comprises attachment of adenosine 5’-*O*-monophosphate (AMP) onto serine, threonine and tyrosine amino acid side chains (**Figure 1A**) (Sieber et al., 2020). Thus far two AMP transferases are known in metazoans to catalyze protein AMPylation, protein adenylyltransferase FICD (FICD) and SelO (SELENOO) (Casey and Orth, 2017; Sreelatha et al., 2018). FICD is characterized by its endoplasmic reticulum (ER) localization and dual catalytical activity of AMPylation and deAMPylation.(Preissler et al., 2017; Sengupta et al., 2019) FICD catalyzes an AMP transfer from its substrate ATP and reverses the modification by the hydrolysis of the AMP-protein ester. FICD’s catalytic activity is regulated by its α-helix inhibition loop through the interaction of Glu234 positioned in the inhibition loop and Arg374, which is necessary for a complexation of ATP in the active site (Engel et al., 2012). Initial biochemical studies identified ER localized heat shock protein HSPA5 (also called BiP or GRP78) as a cognate substrate of FICD (Ham et al., 2014; Sanyal et al., 2015). AMPylation of HSPA5 inhibits its chaperon activity and the downstream unfolded protein response cascade (Preissler et al., 2015). Furthermore, FICD’s activity was recently shown to accelerate the neuronal differentiation of progenitor cells in human cerebral organoids (COs), a tissue model of the human cerebral cortex (Kielkowski et al., 2020a). Surprisingly, apart from the increased number of neurons in tissue overexpressing FICD, some neurons displayed migratory defects. In addition, the FICD was shown to AMPylate α-synuclein *in vitro* (Sanyal et al., 2019). In *Caenorhabditis elegans* FICD has been reported to regulate aggregation of amyloid-β and α-synuclein (Truttmann et al., 2018). In contrast, there are only scare data about the SELO’s function and protein substrates (Sreelatha et al., 2018).

Several complementary strategies were introduced to analyze protein AMPylation, including isotopically labeled adenosine probes (Pieles et al., 2014), radioactively labelled adenosine nucleotides, antibodies (Kingdon et al., 1967; Yarbrough et al., 2009), microarrays (Yu and LaBaer, 2015), *N*^6^-biotin modified ATP (Sreelatha et al., 2018), and *N*^6^-propargyl adenosine 5’-*O*-triphosphate (N6pATP) or phosphoramidate (Broncel et al., 2012; Grammel et al., 2011; Kielkowski et al., 2020b, 2020a). We have recently developed a chemical proteomic approach, which allows high-throughput screening of protein AMPylation and comparison of the AMPylation levels between different conditions. This strategy utilizes the *N*^6^-propargyl or *N*^6^- ethylazide adenosine phosphoramidate (pro-N6pA and pro-N6azA, respectively) probes which upon uptake into cells are metabolically activated to the corresponding *N*^6^-modified adenosine triphosphate and used by endogenous AMP-transferases for protein AMPylation (**Figure 1B**) (Kielkowski et al., 2020b, 2020a). Of note, the resulting *N*^6^-modified ATP is inherently in competition with endogenous ATP and thus cannot report on the exact stoichiometry of protein AMPylation. The application of this chemical proteomic strategy in various cell types has revealed that protein AMPylation is more prevalent than previously appreciated. Interestingly, it has led to the identification of a large group protein substrates from different subcellular compartments not restricted to ER, including cytosolic (PFKP, SQSTM1), nucleolar (PPME1), cytoskeletal (TUBB, MAP2), and lysosomal (CTSB, PLD3, and ACP2) proteins (Broncel et al., 2016; Kielkowski et al., 2020a). In particular, except N-glycosylation there is only limited evidence on PTMs of luminal lysosomal proteins amid numerous diseases associated with lysosomal proteins such as lysosomal acid phosphatase ACP2 in lysosomal storage diseases (LSD) or 5’-3’ exonuclease PLD3 in Alzheimer’s diseases (Cappel et al., 2021; Cruchaga et al., 2014; Gonzalez et al., 2018; Marques and Saftig, 2019; Schultz et al., 2011; Stadlmann et al., 2017; Stoka et al., 2016). Therefore, discovery of lysosomal protein PTMs might shed a new light onto the regulation of their function, localization, and protein-protein interactions and hence putatively uncover unknown pathophysiologicl mechanisms.

**Figure 1.**
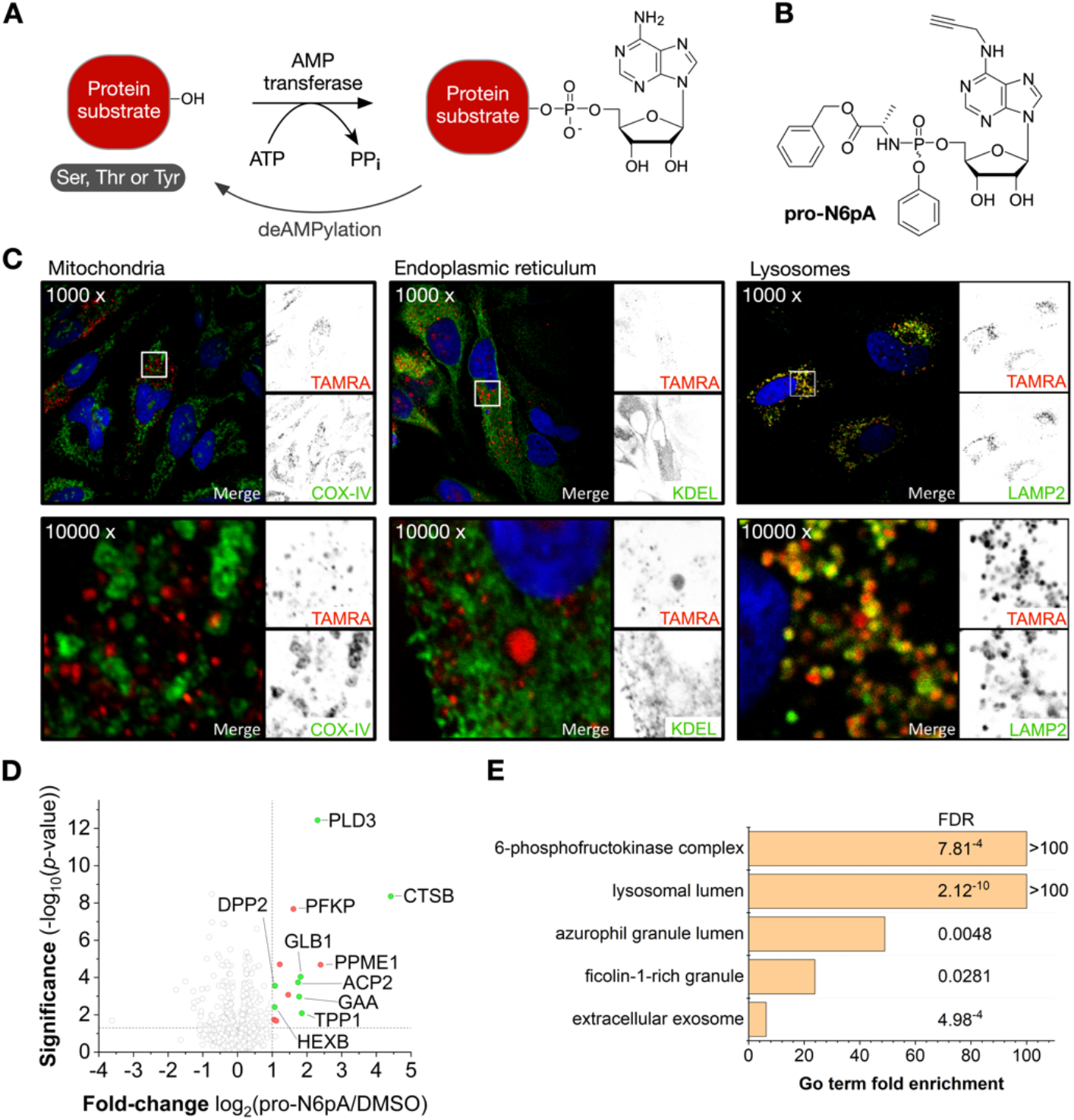
Human neuroblastoma cells (SH-SY5Y) display an enrichment of AMPylated proteins in lysosomes. **A)** Schematic representation of protein AMPylation. **B)** Structure of the pro-N6pA probe for *in situ* labelling of AMPylated proteins. **C)** Click chemistry staining of pro- N6pA with TAMRA-N_3_ (red) and immunocytochemical staining in SH-SY5Y and colocalizations with markers for lysosome (LAMP2), ER (KDEL) and mitochondria (COX-IV). **D)** Volcano plot showing the significantly enriched AMPylated proteins from SH-SY5Y cells. Proteins localized to lysosome are depicted in green. Significantly enriched protein hits with other subcellular localization are in red (cut-off lines at 2-fold enrichment and *p*-value of 0.05). **E)** GO terms analysis of significantly enriched proteins in SH-SY5Y cells.

Here, we characterize protein AMPylation during neuronal differentiation and maturation by a combination of chemical proteomics and gel-based approach showing the two lysosomal proteins PLD3 and ACP2 to be increasingly AMPylated during neural differentiation and in case of PLD3 inhibiting its catalytic activity.

## Results

### AMPylated proteins localize to lysosomes in SH-SY5Y neuroblastoma cells

We have developed a chemical proteomic strategy in previous studies to analyze the protein AMPylation in living cells using the cell-permeable pro-N6pA probe (Kielkowski et al., 2020b, 2020a). In this study, we have started with the completion of this data set with fluorescence imaging of the SH-SY5Y cells treated with the pro-N6pA probe to visualize the subcellular localization of AMPylated proteins. Interestingly, probe-labelled proteins exhibit a vesicular distribution pattern overlapping with lysosomes as confirmed by co-staining with lysosomal specific marker LAMP2 (**Figure 1C**). Colocalization of the majority of AMP-probe signal with the ER and mitochondria was excluded by co-staining of the pro-N6pA probe with antibodies against KDEL and COX-IV, respectively. These findings point to an enrichment of AMPylated proteins in lysosomes and raise questions about the localization and origin of their AMPylation. GO analysis of chemical proteomics data from SH-SY5Y cells indicated an overrepresentation of lysosomal proteins among those enriched for AMPylation by 54 % of all significantly enriched proteins. These include CTSB, PLD3, GAA, GLB1, TPP1, HEXB, DPP2, and GUSB, with PLD3 and CTSB showing the strongest enrichment (**Figure 1D, E**). Next, we asked whether AMPylation of lysosomal proteins is changing during neuronal differentiation and maturation and how the AMPylation status correlates with non-neuronal cell types. Although the chemical proteomic analysis of protein AMPylation variation during neuronal differentiation was previously performed, it has focused solely on two differentiation stages, neuronal progenitors (NPCs) and mature neurons. To investigate the AMPylation of lysosomal proteins in differentiating neurons with higher temporal resolution, we searched for a suitable cellular system, which would allow collecting sufficient amounts of total protein amounts for chemical proteomic analysis in shorter periods of time, thus overcoming the bottlenecks of the standard human-induced pluripotent stem cells (iPSCs) differentiation protocols (Boyer et al., 2012).

### Chemical proteomic analysis of protein AMPylation in differentiating iNGN cells

To map the changes in protein AMPylation during the neuronal differentiation in more detail we took advantage of the human-induced pluripotent stem cells with inducible overexpression of a pair of transcription factors, Neurogenin-1 and Neurogenin-2, leading to their rapid differentiation into a homogenous population of functional bipolar neurons within 4 days (iNGN) (Busskamp et al., 2014). The changes in AMPylation were followed at six time points during the iNGN differentiation and maturation using a previously described chemical proteomics strategy utilizing a metabolically activated pro-N6pA probe (**Figure 2A**). In brief, cells are treated with the pro-N6pA probe 16 h prior to their harvest to allow for metabolic activation to corresponding *N*^6^-propargyl ATP and labelling of the proteins by endogenous AMP-transferases. Subsequently, the cells are lysed, and alkyne modified proteins are further coupled with biotin-PEG-azide by Cu(I) catalyzed click chemistry. The probe-AMPylated and biotinylated proteins were then enriched on avidin-coated agarose-beads and on-beads trypsinized to give complex peptides mixtures which were resolved by LC-MS/MS measurement. The resulting label-free quantification (LFQ) of proteins from four replicates and their comparison with DMSO treated cells prepared in the same manner provided the quantitative differences in protein AMPylations between undifferentiated and differentiated iNGN cells (**Figure 2B, Suppelmental Figure 1** and **Supplemental Table 1–6** and **8**). The background from unspecific protein binding to the avidin-agarose-beads in both vehicle control (DMSO) and probe treated cells partially reflects the total protein expression level. To correct for this contribution, we have carried out the whole proteome analysis of iNGNs during the course of differentiation (**Supplemental Figure 2**). Additionally, the whole proteome analysis confirmed the identity of the cells and the progress of neuronal differentiation and maturation (**Supplemental Figure 3** and **Supplementaly Table 7, 8**). The examination of the profile plots of enriched proteins show a distinct pattern of AMPylation dynamics during neuronal maturation (**Figure 2C**). The most distinct composition was observed for the two lysosomal proteins ACP2 and ABHD6, with a linear increase of AMPylation during the course of differentiation, while the total expression level of both proteins remains stable during the process (**Figure 2C** and **Supplemental Figure 2**). On the other hand, other lysosomal AMPylated proteins show a stable AMPylation level, for example, CTSC. Similar, increase during iNGNs differentiation was observed for the cytosolic protein ATP-dependent 6- phosphofructokinase PFKP, a gatekeeper of glycolysis (**Supplemental Figure 2**). Next, we focused on the lysosomal protein PLD3, which was recently associated with Alzheimer’s disease, but the physiological function in neurons and regulation of which has been thus far controversial (Arranz et al., 2017; Cruchaga et al., 2014). As shown previously, whole proteome analysis of PLD3 exhibit an increase of the PLD3 in neurons as compared to the undifferentiated iNGNs. The PLD3 is known to be transported from the ER and the Golgi to lysosomes through endosomes where the cytosolic N-terminal membrane-bound domain of the full-length PLD3 is proteolytically cleaved, yielding the soluble luminal PLD3 containing the putative active site which is delivered to lysosomes (Gonzalez et al., 2018). Both chemical proteomics and whole proteome analysis cannot provide the information as to which form of PLD3 is AMPylated, thus we have turned our focus on the development of a gel-based methodology which would allow for separation of different protein forms as well as to distinguish protein AMPylation. Previously, the isoelectric focusing gels have been used for the separation of AMPylated HSPA5 from its unmodified form, but the separation of the two species appears to be rather limited (Preissler et al., 2015). Therefore, we explored the possibility to use another gel-based method, which also takes advantage of the presence of a phosphate moiety, but utilize the Phos-tag ligand (Kinoshita et al., 2004, 2006).

**Figure 2.**
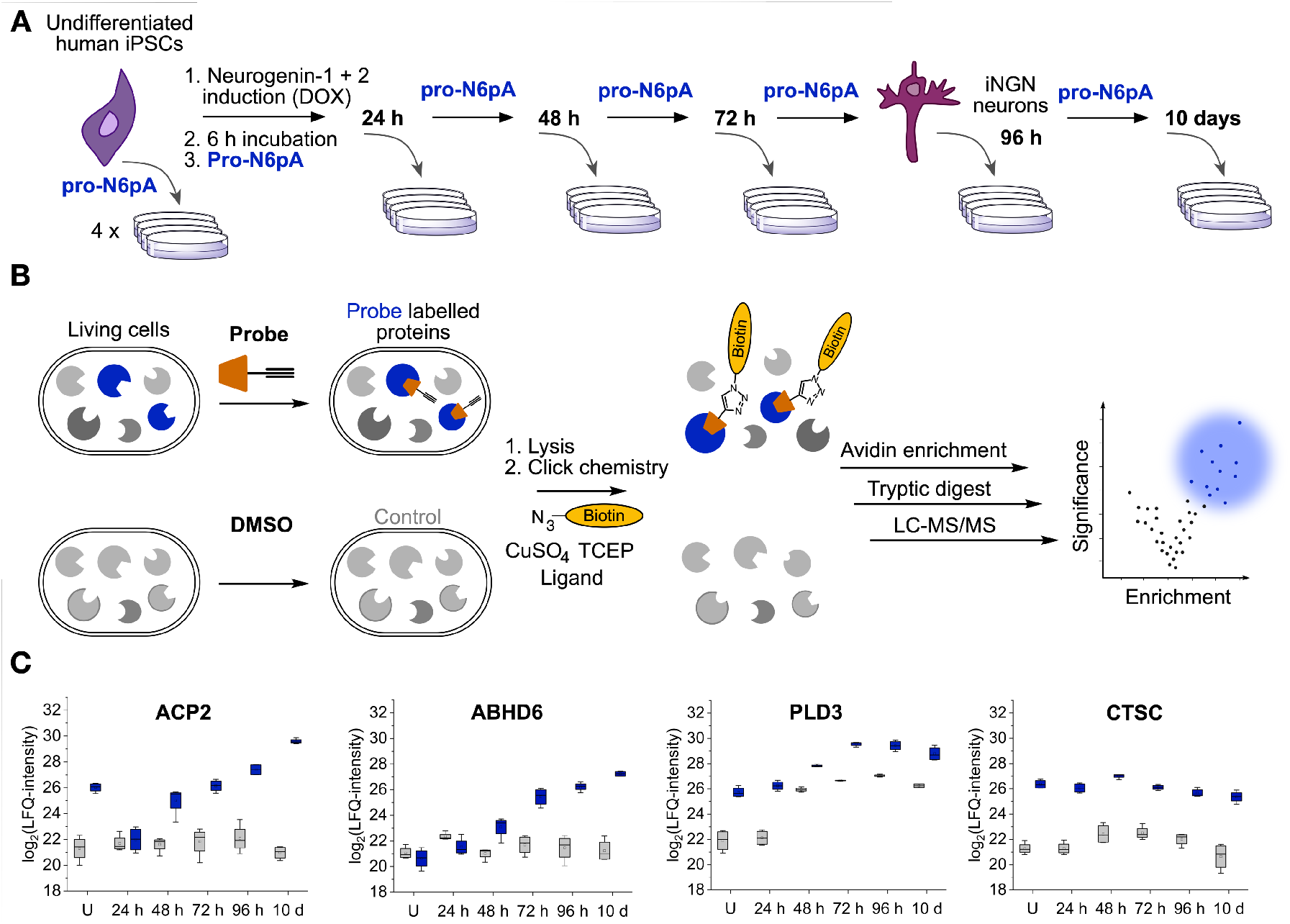
Chemical proteomics of neuronal differentiation and maturation shows specific patterns of protein AMPylation. **A)** Schematic iNGNs differentiation procedure and pro-N6pA probe treatment. **B)** Chemical proteomic protocol comparing the probe and DMSO control treated cells. **C)** Profile plots of significantly enriched proteins ACP2, ABHD6, PLD3 and CTSC from pro-N6pA probe treated iNGNs. Blue boxes represent the LFQ-intensities from pro-N6pA probe treated cells after enrichment. Grey boxes represent LFQ-intensities from control (DMSO treated) cells, showing unspecific binding to avidin-coated agarose-beads (*n* = 4, the box is defined by 25^th^ and 75^th^ percentile, whishers show outliers, line is a median and circle is a mean).

### Phos-tag ligand-based polyacrylamide gel electrophoresis separates the modified and unmodified proteins

In order to validate our results from chemical proteomics experiments by a complementary method, which would not rely on the pro-N6pA probe and mass spectrometry, we have concentrated on the development of a gel-based approach. Therefore, we have tested the possibility to capture the monophosphate moiety of AMPylated proteins using alkoxide-bridge Mn^2+^ metal complex. The commercially available Mn^2+^ ion binding Phos-tag ligand was used for the preparation of sodium dodecyl sulfate (SDS)-polyacrylamide gels (SDS-PAGE) and for separation of the recombinant Rab1b protein, which was AMPylated *in vitro* by recombinant bacterial effector DrrA (Du et al., 2021). The Rab1b identities were confirmed by the top-down mass spectrometry (**Supplemental Figure 4**). Direct comparison of non-AMPylated, AMPylated and 1:1 mixture of both modified and unmodified Rab1b showed that indeed the Phos-tag ligand added to standard SDS-PAGE resolves the two species and yields two clearly separated bands as visualized by Coomassie staining (**Figure 3A**). A control experiment using the SDS-PAGE without addition of the Phos-tag ligand showed no separation of the two species (**Figure 3A**). Because the Phos-tag ligand was initially developed for separation of the phosphorylated proteins, we have treated the *in vitro* AMPylated Rab1b with shrimp alkaline phosphatase to exclude the potential separation due to protein phosphorylation and to show the resistance of the AMP moiety against the phosphatase cleavage (**Figure 3A**). The activity of the phosphatase was verified by hydrolysis of the phosphorylated ovalbumin (**Supplemental Figure 5**). Next, we examined whether it is possible to separate AMPylated HSPA5 from its unmodified form in HeLa cell lysates. Following the Phos-tag gel separation, the gel was blotted onto a PVDF membrane and visualized by incubation with an anti-HSPA5 antibody. We have observed a clear separation of the two forms confirming our hypothesis that the Phos-tag ligand can be used for separation of the AMPylated proteins (**Supplemental Figure 6**). Thus, we have established the Phos-tag ligand functionalized SDS-PAGE separation as a useful method for analysis of the AMPylated proteins, which overcomes the necessity for treatment with the probe, and can thus be used for analysis of protein AMPylation from wider range of sources, for example from animal tissues. In the chemical proteomics experiment, we have identified several AMPylated lysosomal proteins, including PLD3 and ACP2. To validate these results, we have performed the Phos-tag gel-based separation from undifferentiated iNGNs and one to ten days after doxycycline induction of differentiation. To our surprise, we observed the AMPylated soluble form of PLD3 only after four days of differentiation, with a majority of AMPylated PLD3 in ten days differentiated neurons which is in line with our observation of increasing PLD3 AMPylation from the chemical proteomics study (**Figure 3B** and **Supplemental Figure 7**). Moreover, in case of the PLD3, analysis by standard SDS-PAGE showed that the soluble lysosomal PLD3 form increases drammatically with increasing time of iNGN differentiation (up to 72 h), while it drops substantially upon maturation of iNGN neurons between 4 and 10 days post induction (**Figure 3B**). Additionally, a similar AMPylation trend was observed for another lysosomal protein, ACP2, further corroborating the chemical proteomic results and pointing to a specific function of protein AMPylation in neuronal maturation (**Figure 3B**) (Makrypidi et al., 2012). To exlude the separation due to the protein phosphorylation, lysates from the 10d differentiated iNGNs were treated with the shrimp alkaline phosphatase (**Figure 3C**). Taken together, the combination of standard SDS-PAGE and Phos-tag-based gel electrophoresis enabled detailed characterization of PLD3 post-translational processing and AMPylation dynamics during neuronal differentiation and maturation. The previously described trafficking of the PLD3 demonstrated the processing of the PLD3 by cleavage of its N-terminus containing the N-terminal transmembrane domain (**Figure 3D**) and trafficking to early endosomes and subsequently to lysosomes (Gonzalez et al., 2018). In lysosomes, PLD3 is present as the soluble and active protein form. The amount of the active soluble PLD3 increases during the differentiation process (up to 72), while its AMPylation raises during maturation of iNGN neurons (96 h – 10d). However, the identity and localization of the AMP transferase catalyzing the PLD3 AMPylation remains unknown, thus leaving two plausible scenarios of its AMPylation, processing and trafficking into lysosomes (**Figure 3E**).

**Figure 3.**
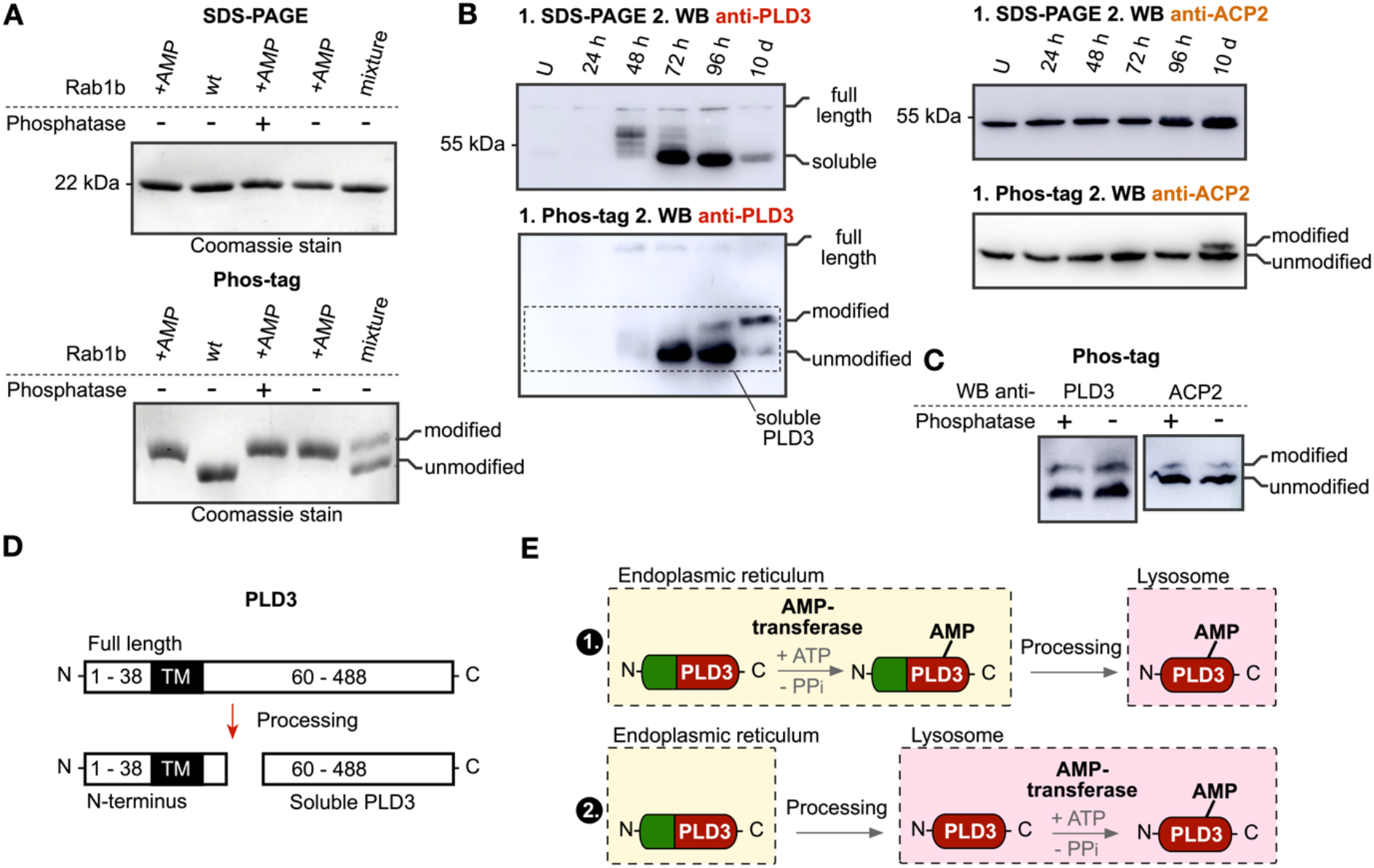
Phos-tag ligand SDS-PAGE distinguishes unmodified and AMPylated proteins. **A)** Coomassie-stained SDS-PAGE and Phos-tag ligand containing SDS-PAGE gel separation of unmodified (wt) and AMPylated (+AMP) recombinant Rab1b. With and without treatment with the shrimp alkaline phosphatase (±). **B)** PLD3 and ACP2 Phos-tag ligand SDS-PAGE separation and Western bloting during iNGNs differentiation and maturation. Visualization using the anti-PLD3 and anti-ACP2 antibodies. **C)** Phos-tag SDS-PAGE analysis of the PLD3 and ACP2 AMPylation after phosphatase treatment. **D**) Processing of the full length PLD3 into soluble active PLD3. **E)** Scheme showing the putative intracellular trafficking and AMPylation of the PLD3.

### AMPylation inhibits the PLD3 activity in iNGN neurons

To correlate catalytical activity of PLD3 with its AMPylation status during the course of iNGNs differentiation a PLD3-specific acid 5’ exonuclease activity assay was carried out (Cappel et al., 2021). Therefore, whole cell lysates of iNGN cells were incubated with a fluorophore- and quencher-coupled oligodesoxynucleotide. 5’ exonucleolytic digest thereby led to a proportional increase in fluorescence signal, measured kinetically over 12 h. The overall activity of the PLD3 per total cellular protein increases during the differentiation which can be accounted for by the increasing levels of its soluble form in lysosomes. In contrast, the PLD3 activity decreases significantly from the day fourth to the tenth day after induction of the differentiation, coinciding with increasing AMPylation of the soluble form during neuronal maturation (**Figure 4A**). This observation suggests that AMPylation inhibits PLD3’s catalytic activity, which is in line with previous reports on inhibition of the chaperon activity of HSPA5 and the peptidase activity of CTSB.

**Figure 4.**
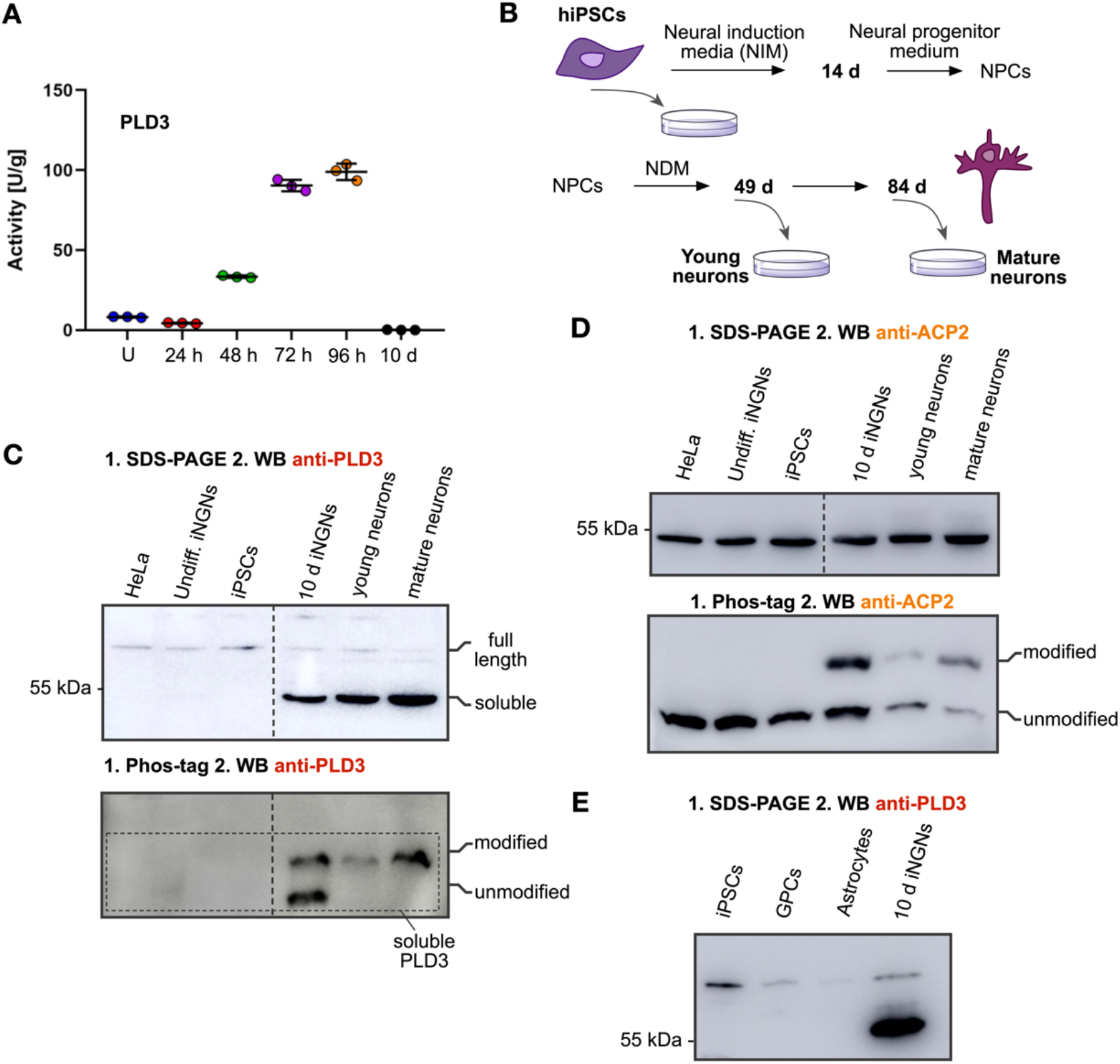
AMPylation inhibits PLD3 activity in neurons. **A)** PLD3 5’ exonuclease activity assay correlates the amount of the soluble PLD3 form with its activity and shows the AMPylation to inhibit the catalytic activity. **B)** Directed differentiation protocol of human iPSCs into dopaminergic neurons. NDM: neural differentiation medium. **C)** SDS-PAGE and Phos-tag ligand SDS-PAGE separations followed by Western blot using the anti-PLD3 antibody shows pronounced and nearly quantitative AMPylation of PLD3 in dopaminergic neurons compared to the iNGN forward reprogramming. **D)** ACP2 SDS-PAGE and Phos-tag ligand SDS-PAGE separations followed by Western blot using anti-ACP2 antibody. **E)** SDS-PAGE followed by Western blot using the anti-PLD3 antibody detects solely the full-length PLD3 in iPSCs, GPCs and astrocytes.

### Directed differentiation of physiological human iPSCs shows the AMPylated PLD3 as the only soluble form in young and mature neurons

In order to compare the AMPylation pattern obtained from the highly homogenous and robust differentiation and maturation of iNGN cells we have collected lysates from physiological iPSCs, young and mature neurons differentiated for five and ten weeks respectively (**Figure 4B** and **Supplemental Figure 8**) (Gunhanlar 2018). The AMPylated soluble form was the only PLD3 species detectable by Phos-tag SDS-PAGE in young as well as in mature neurons, while in undifferentiated iPSCs the anti-PLD3 antibody detected no soluble form, whether unmodified or AMPylated (**Figure 4C** and **Supplemental Figure 9**). Comparison of PLD3 pattern in ten days differentiated iNGNs, young and mature physiological neurons with that in lysates from HeLa cells and undifferentiated iPSCs and iNGNs shows that the soluble forms, whether unmodified or AMPylated, are specific to neurons. Moreover, the analysis of PLD3 processing and AMPylation during differentiation of iPSCs into glial progenitor cells (GPCs) and astrocytes confirmed the specificity of this process for neuronal cell lineage (**Figure 4E**, see **Supplemental Figure 9** for Phos-tag SDS-PAGE analysis). A similar degree of ACP2 AMPylation was seen in ten days differentiated iNGN and physiological mature neural networks (**Figure 4D**). Taken together, these findings further corroborate the hypothesis that AMPylation plays a specific role in maturation of human neurons.

## Discussion

Lysosomal dysfunction is linked to several human pathologies such as LSD’s, cancer, neurodegeneration, and ageing (Marques and Saftig, 2019). Neuronal cells are particularly sensitive to impaired lysosomal function due to their tight control differentiation process and postmitotic character. Even though the genetic basis and the biochemistry underlying these diseases, are known the cellular and molecular mechanisms leading to disruption of neuronal viability, remains to be understood. Our study describes lysosomal proteins to be significantly enriched amongst the AMPylated proteins with changing stoichiometry at various stages of neuronal differentiation, pointing to a specific function of protein AMPylation during differentiation and maturation. The fine tunning of the lysosomal activity was recently reported to be critical for maintenance of the quiescent neural stem cells fitness and their activation responsiveness. Thus, the precise orchestration of these processes might be achieved by adding an extra regulatory layer of protein PTMs, including AMPylation (Leeman 2018). Furthermore, comparing the expression levels of identified AMPylated proteins during development of mouse and human embryos as well as human organoids show cleare differences between the species and model system pointing to regulation of the proteins at different levels and stages of gestation (**Supplemental Figure 10**) (Klingler 2021). The gel-based analyses of PLD3 and ACP2 showed a striking increase of the AMPylated soluble form in differentiated iNGN neurons. Comparing the iNGN forward reprogramming model of neuronal differentiation with differentiation of physiological human iPSCs into young and mature neurons showed an even stronger trend with fully modified PLD3 in mature physiological neurons. Prologing the neurogenic period give rise to more neurons and expansion of the upper-layer of neocortex (Stepien 2020). Thus, lengthening the maturation period of the physiological neurons in comparison to the fast differentiation of iNGN cells may provide more time to established more strictly defined AMPylation status of PLD3. In particular, the increased expression of the PLD3 was shown to coincide with late neuronal development in the hippocampus and the primary somatosensory cortex (Pedersen 1998). In consequence, the PLD3 AMPylation may modulate PLD3 activity or stability to ensure the proper lysosomal function during the migration and maturation of the basal progenitors from the cortical subventricular zone. The activity assay of PLD3 revealed the strong inhibition of PLD3’s activity upon AMPylation in iNGN neurons. The observation is in strong correlation with the levels of the lysosomal PLD3 soluble form and the AMPylation status. Even though we have characterized in detail the progress of PLD3 AMPylation, it remains to be elucidated where the AMPylation of the PLD3 occurs and which AMP-transferase catalyzes the attachment of this modification (**Figure 3E**) or if it can be reversed by active deAMPylation. The heterogeneity of the cell types in the human cortex is enormouse including excitatory and inhibitory neurons, astrocytes, microglia and oligodendrocytes. Analysis of the expression levels of the lysosomal proteins ACP2, ABHD6, PLD3 and CTSC in these cell types exhibit the substantial differences (**Supplemental Figure 11**) (Kanton 2019). Together with considerable variance in the AMPylation status, as we have demonstrated for PLD3 in astrocytes and neuronal cell lineage, it highlights the diversity and specificity of the proteoforms in central nervous system. Therefore, in future studies, it will be important to compare the AMPylation status of these proteins in the different cell types and to elucidate the functional cosequences. Our study uncovers the rare modification of the lysosomal proteins and thus provides the insight into the mechanism modulating the protein activity in this critical cellular department. This finding may open up a new possibilities for designing novel therapeutically strategies against the damage linked to lysosomal dysfunction during neurodevelopment, cancer and ageing.

## Supporting information

Supplemental Resource Table

## Author contributions

P.K., M.D. and T.B. have conceived the study. T.B. has cultured and differentiated the iNGN cells. T.B. has carried out all proteomic experiments and gel-based analyses. C. C. has performed the fluorescence imaging and PLD3 activity assay. F.M. cultured the physiological hiPSCs and differentiated them into neurons. G.S. cultured the hiPSCs and differentiated them into GPCs and astrocytes. E. K. and F.S. have helped to establish the iNGN cell culture and reviewed the manuscript. P.K. and T.B. wrote the manuscript. All authors have revised the manuscript.

## Acknowledgements

This research was supported by the Liebig fellowship from Fonds der chemischen Industrie to P.K. and T.B. We are grateful for the kind gift of recombinant Rab1b wt and AMPylated form from group of Dr. Sabine Schneider by Dr. Marie-Kristin von Wrisberg.

