## Supplemental Figures for "AMPylation is a specific lysosomal protein posttranslational modification in neuronal maturation"

Becker et al.

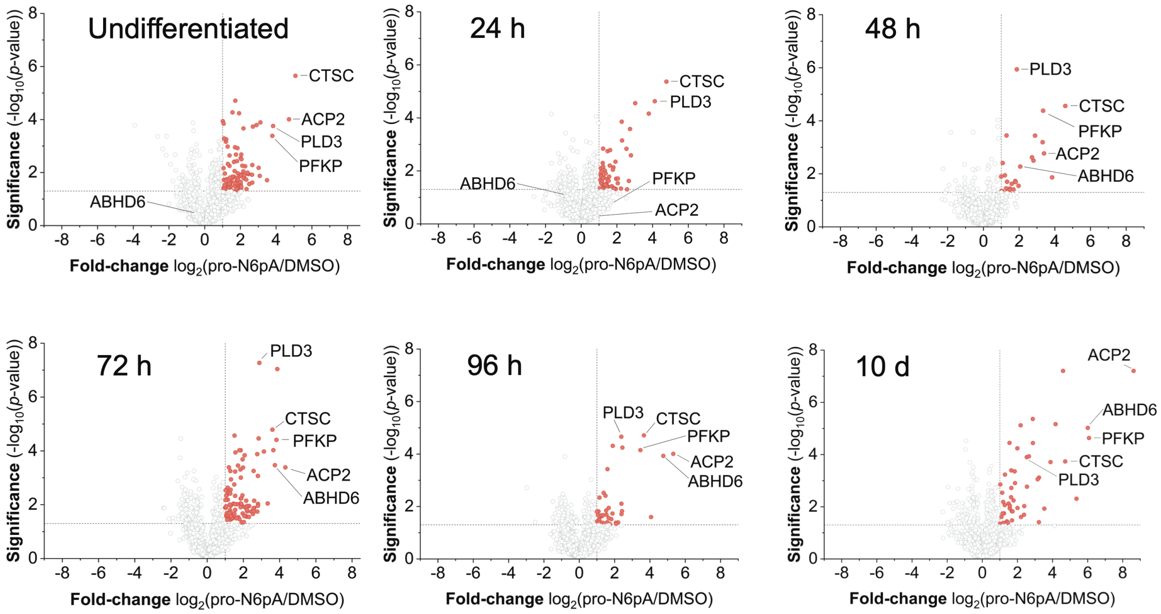


**Supplemental Figure 1.** Volcano plots showing the enrichment of AMPylated proteins at different times during differentiation. Red circles highlight the significantly enriched proteins. ($n$ = 4, cut-off lines are at $p$-value >0.05 and at least 2-fold enrichment).

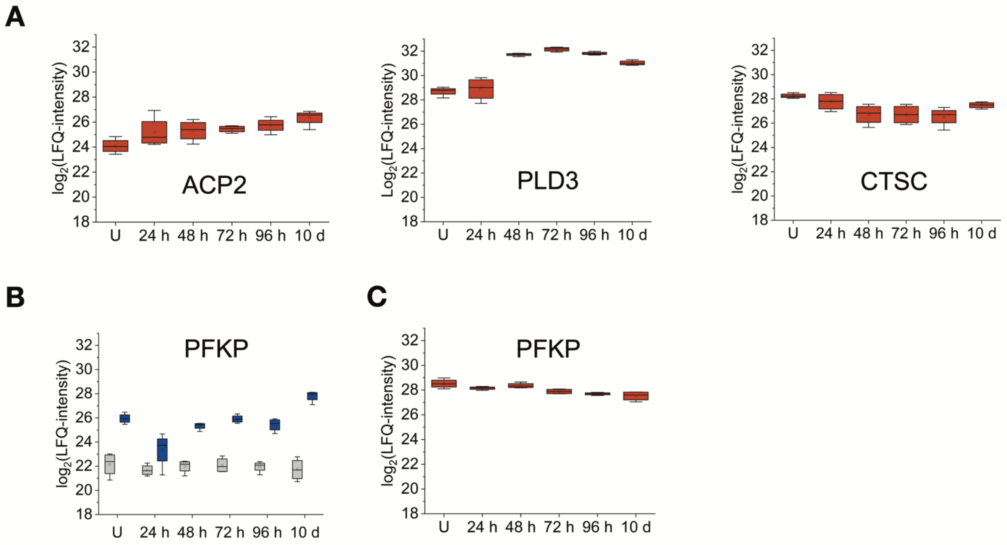


**Supplemental Figure 2. A**) Profile plots showing total protein expressions estimated from the whole proteome analysis during the iNGN differentiation and maturation. **B**) Profile plot visualizing the enrichment (blue boxes) of the PFKP in the chemical proteomic experiment. Grey boxes stand for PFKP background binding to the agarose-beads found in the DMSO treated control. **C**) Profile plot showing total expression of PFKP estimated from the whole proteome analysis during the course of differentiation.

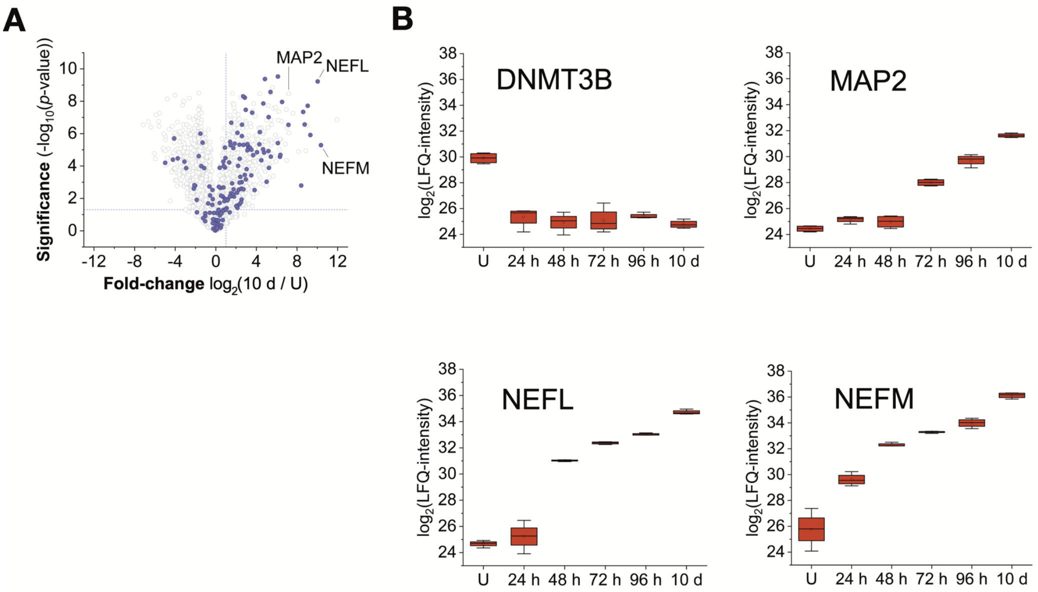


**Supplemental Figure 3.** iNGN differentiation and maturation. **A**) Volcano plot representing the difference in protein expression between undifferentiated iNGNs and 10 d old neurons. Purple circles label protein annotated as neuronal processes by GO terms. B) Selected protein markers of neural maturation.

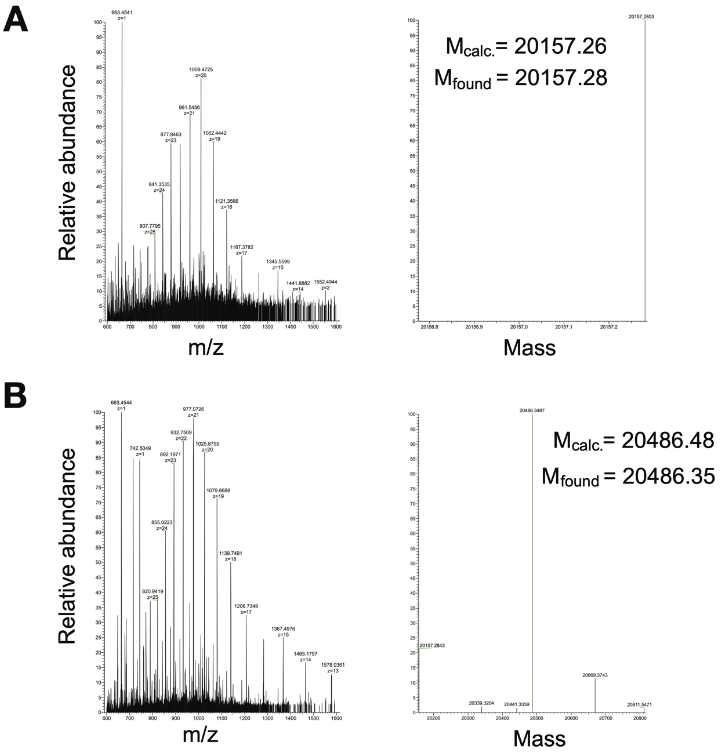


**Supplemental Figure 4.** Top-down mass spectrometry of unmodified (**A**) and AMPylated Rab1b (**B**). ΔM is 329.07 Da corresponding to the AMP moiety.

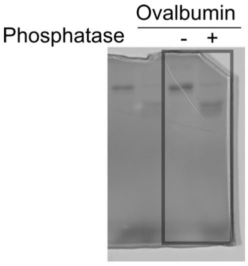


**Supplemental Figure 5.** Phos-tag gel separation of Ovalbumin with and without the phosphatase treatment.

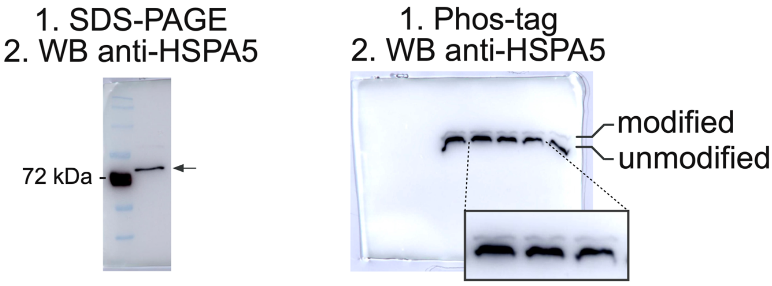


**Supplemental Figure 6.** HSPA5 separation. All lines are from HeLa lysates.

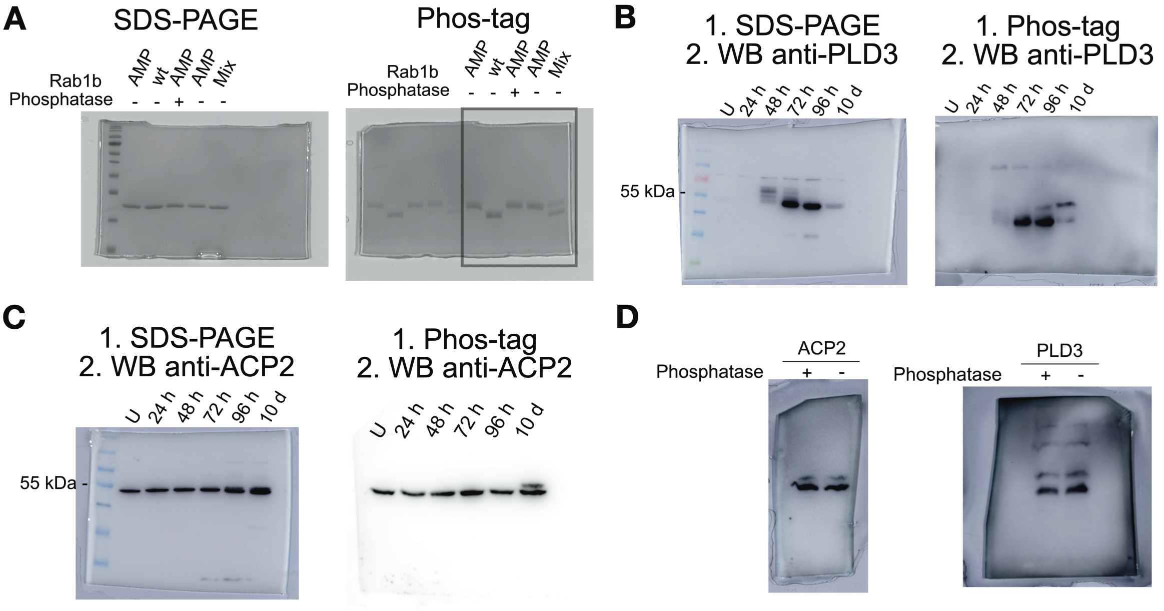


**Supplemental Figure 7.** Uncut SDS-PAGE and Phos-tag gels used in Figure 3. **.A)** Coomassie stained SDS-PAGE and Phos-tag gel separation of AMPylated and non-modified Rab1b. **B**) Western blots of the SDS-PAGE and Phos-tag gel separation of PLD3 during differentiation course of iNGN cells. **C**) Western blots of the SDS-PAGE and Phos-tag gel separation of ACP2 during differentiation course of iNGN cells. **D**) Western blots of the SDS-PAGE and Phos-tag gel separation of PLD3 and ACP2 in 10 days differentiated iNGN cells treated with shrimp alkaline phosphatase.

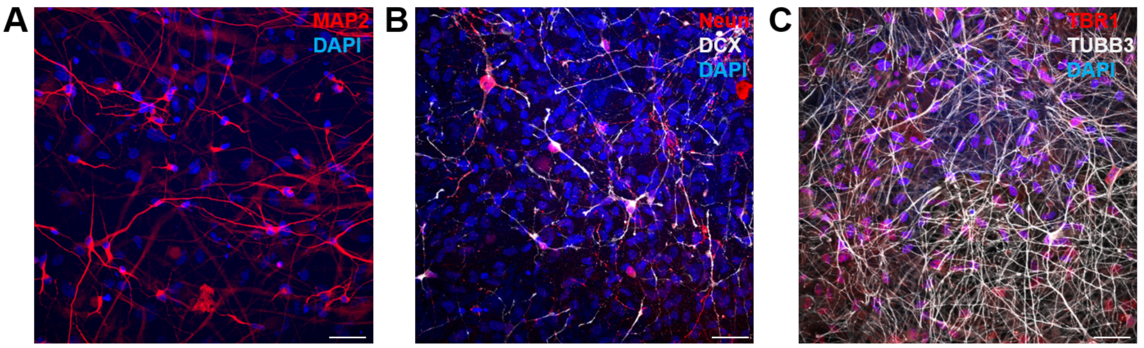


**Supplemental Figure 8.** Characterization of differentiation and maturation of physiological neurons. **A - C**) Micrographs of 2D neuronal cultures after 10 weeks of differentiation. Cells were immunostained for **A**) MAP2, **B**) Neun, Doublecortin (DCX), **C**) TBR1 and TUBB3. Nuclei (blue) are stained with DAPI. Scale bars: 50 μm.

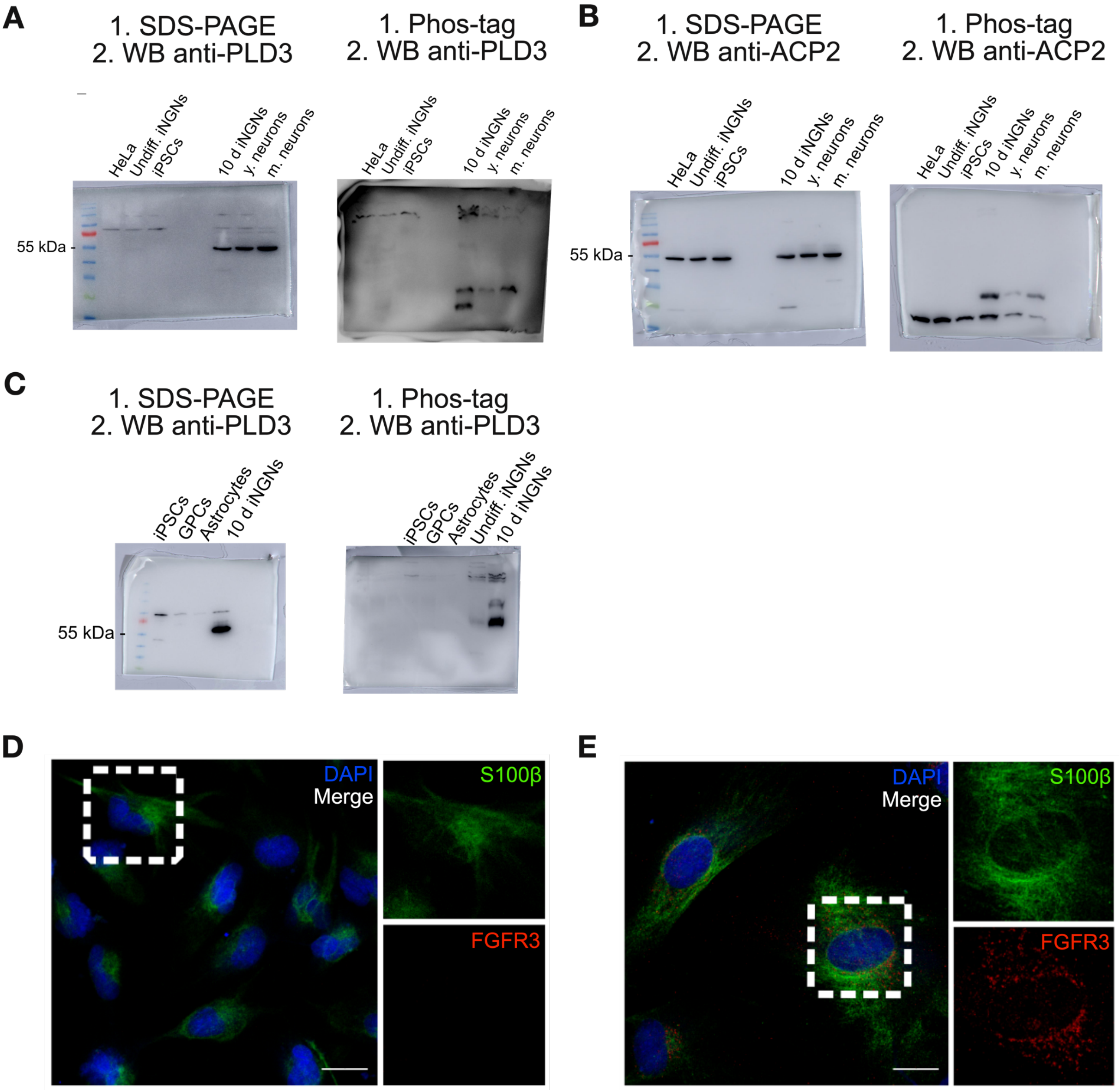


**Supplemental Figure 9.** Uncut SDS-PAGE and Phos-tag gels used in Figure 4. **A**) Western blots of the SDS-PAGE and Phos-tag gel separation of PLD3 comparing HeLa, iNGNs and dopaminergic neurons. **B**) Western blots of the SDS-PAGE and Phos-tag gel separation of ACP2 comparing HeLa, iNGNs and dopaminergic neurons. **D**) and **E**) Characterization of astrocytes differentiation. Nuclei (blue) are stained with DAPI. Scale bars: 20 µm. **D**) GPCs. **E**) Astrocytes.

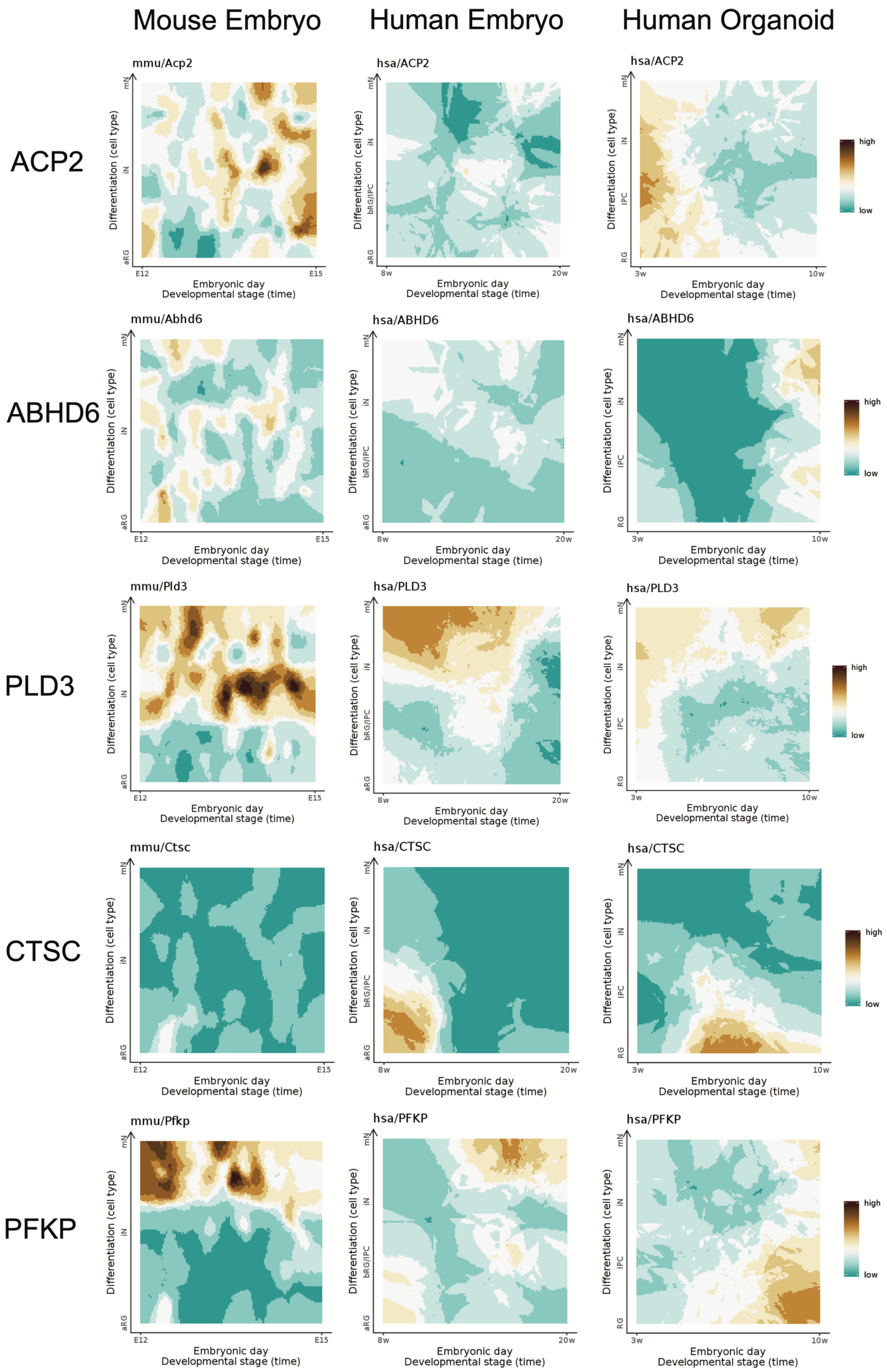


**Supplemental Figure 10.** Protein expression changes during embryo development.

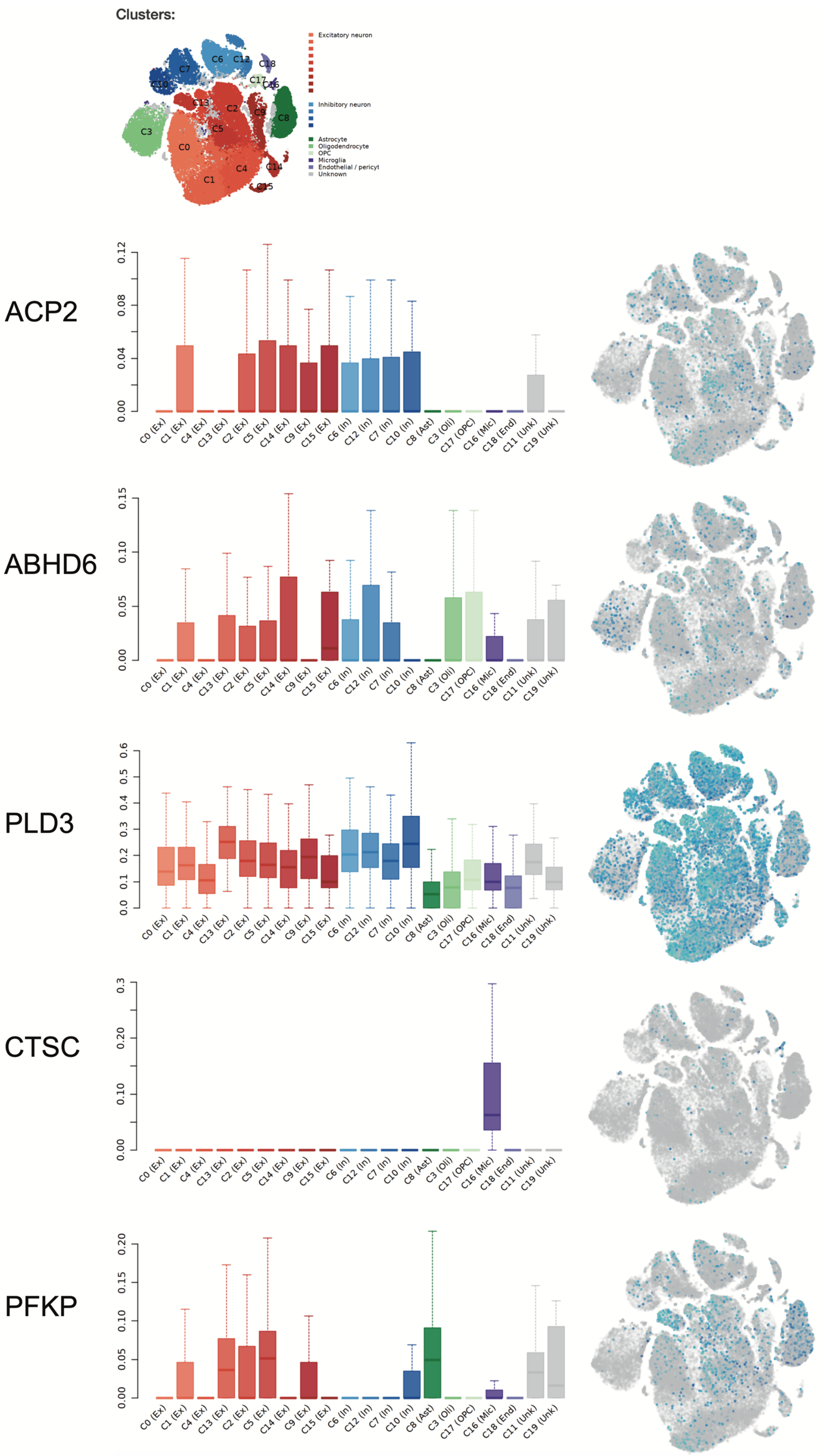


**Supplemental Figure 11.** Analysis of protein expression levels in various cell types.
