## Supplemental Information for "AMPylation is a specific lysosomal protein posttranslational modification in neuronal maturation"

Becker et al.

**Supplemental Information**

***iNGN cells***

***Coating of petri dishes for iNGN culturing***

In order to culture iNGNs, petri dishes have to be coated. For this, 15 mg/mL Geltrex LDEV-free was diluted 1:1000 in cold coating media (49.5 % DMEM (1x), 49.5% F-12, 1 % Pen-Strep 100x) and was immediately added onto the petri dish. The coating volumes were as follows: p60, 3.5 mL; p100, 10 mL and p150, 25 mL. After addition of the coating media to the petri dishes, they were incubated at least one hour in the incubator at 37 °C and were then ready for use.

***Passaging and culturing of iNGN cells in p100 dish***

First, media was removed and cells were washed with 4 mL PBS. Then, 1.5 mL TrypLE^TM^ Express was added and the mixture was incubated 7 min at 37 °C. The detached cells were resuspended with 2 mL pre-warmed E7 media (49 % (v/v) DMEM, 49 % (v/v) F-12, 1 % (v/v) **Alanyl-Glutamine** 100x, 77.6 nM Na_2_SeO_3_, 11.2 mM NaCl, 10 µg/mL hHolo-Transferrin, 10 µg/mL hInsulin) supplemented with 2 µM thiazovivin (E7+TZ) and the cell suspension was transferred to a 15 mL falcon. Before the cell suspension was spinned down 5 min at 600 rpm, the concentration of the cell suspension was determined by mixing 10 µL cell suspension with 10 µL trypan blue. Afterwards, the supernatant was removed and the cell pellet was resuspended in the calculated amount of media to obtain a concentration of 1.5 million cells in 1 mL (For seeding iNGNs for differentiation the concentration was adjusted to 2.5 million cells in 1 mL). Next, the media of the previously coated p100 dish was removed and 9 mL E7+TZ media was added and topped with 1 mL of the cell suspension. Finally, the E7+TZ media was supplemented with 10 µL TGF-beta (2.0 µg/mL) and 10 µL FGF-IS (20 µg/mL) in order to obtain E9+TZ medium. The iNGN cells were cultured overnight at 37 °C and 5 % CO_2_, before the media was exchanged to E9 on the next day.

***Differentiation of iNGNs in p100 dish***

In order to differentiate iNGNs to neurons during 4 days, the expression of the two transcriptions factors Neurogenin-1 and Neurogenin-2 have to be induced by the addition of doxycycline (Dox). For this, 2.5 million cells in 10 mL E7 media containing 0.5 µg/mL Dox and 2 µM TZ (E7+Dox+TZ) were seeded in a p100 dish. After overnight incubation at 37 °C in the incubator, the media was exchanged to 10 mL E7+Dox without TZ. On the next day, 10 mL fresh E7+Dox media was added onto the dish. On day 4, half of the E7+dox media was removed and 5 mL Neurobasal A media supplemented with 4 % NS-21 was added. On day 5, the complete media was removed and 10 mL Neurobasal A media supplemented with 2 % NS-21 and 2 mM L-alanyl-glutmanine was added. On day 7, 2 mL of the media was removed and 2 mL Neurobasal A media supplemented with 2 % NS-21 and 2 mM L-alanyl-glutmanine was added. Until day 10, the iNGNs were incubated at 37 °C in the incubator without any further media exchange.

***iPSC Culture for physiological neurons differentiation***

iPSCs (RRID: CVCL_YT30) were cultured as previously described in (Ayo-Martin et al., 2020). They were cultured in Matrigel® Basement Membrane Matrix, LDEV-free coated plates in mTESR1 medium supplemented with 1× mTESR1 supplement. Media was changed every day. For passaging, the cells were dissociated using Accutase and the collected colonies were resuspended in mTESR1 with 1× mTESR1 supplement and 10 μM Rock inhibitor Y-27632(2HCl) and diluted in the desired density.

***Generation of NPCs***

Neural progenitors were generated as previously described with modifications. (Boyer et al., 2012; Klaus et al., 2019) In short, embryoid bodies were generated from iPSCs by plating colonies in suspension in neural induction medium consisting of DMEM F12 with N2 and B27 supplements (minus vitamin A). After 7 days in suspension, Embryoid bodies were plated on polyornithine and laminin coated dishes and cultured for 7 days in neural induction medium. Neural rosettes were manually picked, dissociated and plated in a new polyornithine/laminin-coated plate in neural progenitor medium (neural induction medium supplemented with bFGF at 20 ng/ml). For passaging, the cells were dissociated using Accutase and split at a maximum ratio of 1:4.

***Generation of Neurons***

Neurons were generated following the Gunhanlar protocol. (Gunhanlar et al., 2018) Briefly, NPCs were plated on poly-L-ornithine and laminin coated dishes in neural differentiation medium consisting of Neurobasal with N2, B27 supplements (minus vitamin A), minimum essential medium/non-essential amino acid and laminin, supplemented with BDNF, GDNF, ascorbic acid and dcAMP. Media was changed every 2-3 days. Young and mature neurons were collected after 5 weeks and 10 weeks in culture respectively.

***iPSC Culture for Astrocytes differentiation***

iPSCs were cultured on Geltrex™ LDEV-Free, Reduced Growth Factor Basement Membrane Matrix coated 6-well plates in mTESR1 medium supplemented with 1× mTESR1 supplement. Media was changed every day. For passaging, the cells were incubated with Collagenase Type IV for 5–7 minutes at 37°C. The collagenase was aspirated and fresh mTESR1 with 1× mTESR1 supplement was added to each well. A cell scraper was used to collect the cells, and they were subsequently plated on a fresh 6-well plate at the desired dilution.

***Generation of GPCs and Astrocytes***

Glial progenitor cells and astrocytes were generated as previously described with modifications (Santos et al., 2017). Briefly, confluent iPSC cultures were dissociated with collagenase, collected with a cell scraper and then cultured in suspension to form embryoid bodies. The first 24hrs the cells were cultured in mTESR1 with 1× mTESR1 supplement and 10 μM Rock inhibitor Y-27632. For the next two weeks the cells were cultured in Astrocyte medium (AM) supplemented with 20ng/ml Noggin and 10 ng/mL PDGFAA, and an additional week with only PDGFAA. The embryoid bodies were then manually dissociated by pipetting and the resulting GPCs were plated on poly-L-ornithine and laminin-coated dishes in AM supplemented with 10ng/ml bFGF and 10ng/ml EGF. Astrocytes were differentiated from GPCs in AM supplemented with 10ng/ml LIF. Media was changed every other day. The GPCs and astrocytes were collected after 5 weeks and 9 weeks of differentiation respectively. The images were obtained on a LSM710 laser-scanning confocal (Carl Zeiss microscope, ZEN software) at x40 magnification.

***Culturing of HeLa and SH-SY5Y cells***

HeLa (RRID: CVCL_0030) and SH-SY5Y (RRID: CVCL_0019) cells were cultured in Dulbeccos Modified Eagles Medium – high glucose (DMEM) supplemented with 10% fetal calf serum (FCS) and 2 mM L-alanyl-glutamine at 37 °C and 5 % CO_2_ atmosphere.

***Fluorescence imaging***

SH-SY5Y neuroblastoma cells were seeded on glass coverslips and grown in DMEM supplemented with 10 % FCS. For Click staining of AMPylated proteins, cells were treated with 100 µM pro-N6pA for 24 h. After washing twice in cold PBS, the cells were fixed with 4 % paraformaldehyde (PFA) in PBS for 20 min, washed two more times in PBS and permeabilized by incubation with 0,1 % Triton-X-100 in PBS for 5 min. The metabolically labelled proteins were coupled to TAMRA via CuAAC using 1 mM CuSO_4_, 5 mM THPTA, 10 µM TAMRA-PEG-N_3_ and 100 mM sodium ascorbate in PBS for 1 h. Following washing twice, a second fixation in 4 % PFA for 10 min and two more washing steps. For immunostaining, the cells were permeabilized in 0.2 % saponin in PBS for 5 min, followed by quenching with 0.12 % glycine and 0.2 % saponin for 10 min and incubation in blocking solution for 1 h (10 % FCS and 0.2 % saponin in PBS). Primary antibodies were diluted in blocking solution and incubated with the coverslips overnight at 4 °C. After four washing steps in 0.2 % saponin, the cells were incubated with AlexaFluor488 coupled secondary antibodies for 1 h at room temperature, washed four times in 0.2 % saponin and twice in H_2_O. The coverslips were finally placed in 15 µl mounting medium (167 mg/ml Mowiol® 4 – 88, 3% glycerol, 20 mg/ml DABCO®, 1 µg/ml DAPI in PBS) Microscopic images were recorded at an Olympus FV1000 confocal laser scanning microscope using a U Plan S-Apo 100x oil objective (1.40 NA).

***Immunoistochemistry – physiological neurons***

10 weeks old mature neurons were fixed using 4% PFA for 10 min and permeabilized with 0.3% Triton for 5 min. After fixation and permeabilization, cells were blocked with 0.1% Tween, 10% Normal Goat Serum. Primary and secondary antibodies were diluted in blocking solution. Nuclei were visualized using 0.5 mg/ml 4,6‐diamidino‐2‐phenylindole (DAPI). Stained cells were analyzed using a Leica laser‐scanning microscope.

***PLD3 activity assay***

For quantitative determination of PLD3 5’exonuclease activity, lysates were prepared in TRIS-lysis buffer (TBS with 1% (v/v) Triton-X-100 and 1 tablet cOmplete^®^ EDTA-free protease inhibitor cocktail). After collecting the whole cell lysates as described above, lysates were diluted in a final volume of 100 µl MES reaction buffer (50 mM MES, 200 mM NaCl) to a final concentration of 50 ng/µl in a lumox® multiwell 96 plate (Sarstedt). The reaction was started by addition of 100 pmol quenched FAM-ssDNA substrate (6-FAM-ACCATGACGTTC*C*T*G*-BMN-Q535 (Biomers.net) with * indicating a phosphothioate bond). After a pre-incubation period of 30 min fluorescence emission at 528 nm (following excitation at 485 nm) was measured in a microwell plate reader (SynergyHT from BioTek) from below the wells over a period of 12 h every 5 min while incubation at 37 °C. For evaluation, a substrate control without lysate and a lysate control without substrate were measured together with the samples. The 5’exonuclease activity was calculated as the slope of the measured fluorescence in the samples minus both controls.

***Chemical proteomics***

***Seeding of iNGNs for differentiation***

The identification of the AMPylation targets during neuronal differentiation was performed with four controls and four probe treated samples for each time point. In each 10 cm dish 2.5 million iNGNs in 10 mL E7+Dox+TZ media were seeded for differentiation. The media composition that is used for each day of differentiation is described in the section above, differentiation of iNGNs. One day before the cells were harvested, the samples were treated with 5 µL of 100 mM AMPylation probe (pro-N6pA) and the controls with 5 µL DMSO.

***Harvesting and cell lysis***

Cells were washed twice with 2 mL PBS. Then, 500 µL lysis buffer (PBS with 1 % (v/v) NP40 and 1 % (w/v) sodium deoxycholate and 1 tablet protease inhibitors per 10 ml buffer)) was added and cells were scrapped into an Eppendorf tube. The cell suspension was incubated 15 min at 4 °C with agitation, before cells were spinned down 10 min at 12,000 rpm and 4 °C. Subsequently, the cytosolic fraction of the lysate was transferred into a new 2 mL Eppendorf tube.

***Measurement of protein concentrations***

In order to measure the protein concentrations of the lysates bicinchoninic acid assay was performed. First, bovine serum albumin (BSA) standards with concentrations of 12.5, 25, 50, 100, 200 and 400 µg/mL were prepared and samples as well as controls were diluted 40 times to a total volume of 200 µL. To measure standards, samples and controls in triplicates, 50 µL of each was added to three wells of a transparent 96-well plate with flat bottom. Afterwards, 100 µL working reagent (2 µL R2 and 98 µL R1) was added to each well by a multistepper and the plate was incubated 15 min at 60 °C. Then, the absorbance at 620 nm was measured by Tecan and the protein concentrations were calculated. For each replicate, except for the 24 h time point (250 µg), 400 µg of protein was used and the volumes were adjusted to a total volume of 970 µL with 0.2 % SDS in PBS.

***Coupling of AMPylated proteins***

In order to couple the alkyne residue of AMPylated target proteins with Biotin-PEG_3_-N_3_, CuAAC was used. For this reaction, 10 µL 10 mM Biotin-PEG_3_-N_3_, 10 µL 100 mM TCEP and 1.2 µL 83.5 mM TBTA were added to 970 µL of each lysate. The mixture was vortexed and spinned down before 20 µL 50 mM CuSO_4_ was added to initiate the reaction. Finally, the reaction mixture was incubated 1.5 h shaking at 600 rpm and 25 °C in the dark.

***Protein precipitation***

First, each reaction mixture of the previously performed click reaction was transferred to a 15 mL falcon. Then, 4 mL acetone was added to each falcon in order to precipitate the proteins. After 1 h of incubation at -20 °C, proteins were spinned down 15 min at 11,000 rpm and 4 °C. Supernatant was discarded and each pellet was resuspended in 1 mL methanol by sonicating 3 times for 5 s at 20 % intensity. Subsequently, each suspension was transferred to a 1.5 mL Eppendorf tube and was centrifuged 10 min at 11,000 rpm and 4 °C. Each pellet was washed again with 1 mL methanol before it was dissolved in 1 mL 0.2 % SDS in PBS by sonicating 3 times for 5 s at 20 % intensity.

***Avidin beads enrichment***

In order to enrich the biotinylated proteins, avidin agarose beads (Sigma-Aldrich) were used. First, avidin beads were thawed on ice before 50 µL avidin bead suspension for each sample or control was washed 3 times with 1 mL 0.2 % SDS in PBS to equilibrate the beads. After each addition of washing solution, the Eppendorf tube was carefully inverted 10 times, the suspension was centrifuged 2 min at 2000 rpm at room temperature and the supernatant was discarded. Subsequently, each dissolved protein pellet was spinned down at maximum speed for 2 min and room temperature before the supernatant was added to the equilibrated beads. After each avidin bead suspension was incubated 1 h under continuous mixing at room temperature, the beads of each sample or control were washed 3 times with 1 mL 0.2 % SDS in PBS, 2 times with 1 mL 6 M urea in H_2_O and 3 times with 1 mL PBS.

***On beads digest of enriched proteins***

In order to prepare defined peptide fragments for the following MS-measurements, the enriched proteins were digested with trypsin. First, the washed avidin beads were resuspended in 200 µL Xbuffer (7 M urea, 2 M thiourea in 20 mM HEPES pH 7.5). Then, 0.2 µL 1 M DTT was added to reduce the disulfide bonds. Afterwards, each mixture was vortexed and incubated 45 min at room temperature shaking at 600 rpm. Next, 2 µL 550 mM IAA was added to alkylate the cysteine residues. After each mixture was vortexed and incubated 30 min at room temperature shaking at 600 rpm in the dark, 0.8 µL 1 M DTT was added to quench the alkylation. Subsequently, each mixture was vortexed and incubated for another 30 min at room temperature shaking at 600 rpm in the dark before 600 µL 50 mM TEAB was added to increase the pH value to 8. Afterwards, 1.5 µL trypsin (0.5 µg/µL in 50 mM acetic acid) was added to digest the enriched proteins. Finally, each mixture was vortexed and incubated overnight at 37 °C shaking at 600 rpm.

On the next day, 4 µL FA was added to stop the digest. Subsequently, each mixture was vortexed and centrifuged 1 min at 2000 rpm before the supernatant was transferred into a new Eppendorf tube. After 50 µL 0.1 % FA was added to the avidin beads, each mixture was vortexed, the centrifugation step was repeated and the supernatant was again transferred to the new Eppendorf tube. Then, 50 µL 0.1 % FA was added once more to the avidin beads and each mixture was vortexed. Finally, each mixture was centrifuged 3 min at 13000 rpm and the supernatant was transferred to the new Eppendorf tube as before. For the following desalting step of the digested proteins, the pH was checked to be below 3.

***Desalting of peptides mixture***

In order to remove all disturbing salt from the digested proteins, Sep Pak C18 cartridges (50 mg columns, waters) were used. First, cartridges were washed with 1 mL ACN and 1 mL 80 % ACN with 0.5 % FA. Then, cartridges were equilibrated with 3 mL 0.5 % FA before the acidified samples were loaded slowly. Afterwards, cartridges were washed with 3 mL 0.5 % FA. The desalted peptides were eluted 2 times with 250 µL 80% ACN with 0.5 % FA into a LoBind Eppendorf tube. Finally, the eluates were lyophilized.

***Final peptide preparation***

First, 30 µL 1 % FA was added to the lyophilized peptides. Then, the mixture was vortexed, spinned down and sonicated 15 min to dissolve the complete peptides. Afterwards, the dissolved peptides were spinned down and added onto centrifugal filter units. Finally, the solution was centrifuged 1 min at 13000 rpm and the filtrate was transferred into plastic MS vials.

***MS-measurement***

MS-measurements were performed on a Q Exactive HF mass spectrometer (Thermo Fisher Scientific) coupled to an UltiMate^TM^ 3000 Nano HPLC (Thermo Fisher Scientific) via an EASY-Spray^TM^ source (Thermo Fisher Scientific). First, peptides were loaded on a Acclaim PepMap 100 µ-precolumn cartridge (5 µm, 100 Å, 300 µm ID × 5 mm, Thermo Fisher Scientific). Then, peptides were separated at 40 °C on a PicoTip emitter (noncoated, 15 cm, 75 µm ID, 8 µm tip, New Objective) that was in house packed with Reprosil-Pur 120 C18-AQ material (1.9 µm, 120 Å, Dr. A. Maisch GmbH). The gradient was run from 1-36 % ACN supplemented with 0.1% FA during a 120 min method (0-5 min 1 %; 5-8 min to 6 %; 8-98 min to 36 %; 98-100 min to 85 %; 100-105 min wash with 85 %; 105-110 min to 1 %, 110-120 min with 1 %) at a flow rate of 200 nL/min. For measurements of chemical-proteomic samples on Q Exactive HF mass spectrometer, the following settings were used: The Q Exactive HF was operated in dd-MS² mode with the following settings: Polarity: positive; MS^1^ resolution: 120k; MS^1^ AGC target: 3e6 charges; MS^1^ maximum IT: 20 ms; MS1 scan range: m/z 300 – 1750; MS² resolution: 15k; MS² AGC target: 2e5 charges; MS² maximum IT: 100 ms; Top N: 20; isolation window: m/z 1.6; isolation offset: m/z 0.2; HCD stepped normalised collision energy: 28 %; intensity threshold: 5e4 counts; charge exclusion: unassigned, 1, 7, 8, >8; peptide match: off; exclude isotopes: on; dynamic exclusion: 90 s.

***Full proteome analysis***

In order to quantify the proteins levels during neuronal differentiation, a full proteom analysis was performed. For this approach, the same lysates (control samples) that were used for the chemical proteomic experiments were used. First, 100 µg protein of each lysate was diluted to a total volume of 200 µL with 0.2 % SDS in PBS. Then, 800 µL acetone was added and incubated 1 h at – 20 °C to precipitate proteins. Next, proteins were spinned down 15 min at 11000 rpm and 4 °C and the supernatant was discarded. Each pellet was resuspended in 1 mL methanol by sonicating 3 times for 5 s at 20 % intensity and the mixture was spinned down once again as before. The supernatant was discarded and the washed protein pellets were dissolved in 200 µL Xbuffer. The following digest and desalting were performed as described in the sections above. Finally, the lyophilized peptides were dissolved in 200 µL 1 % FA and filtered as described in the section, final peptide preparation.

***Phosphatase treatment***

In order to ensure that we only observed the separation of AMPylated and not phosphorylated proteins using a Phos-tag gel, the lysates were treated with a phosphatase prior to analysis. For the reaction, 10 µL 10x buffer (0.5 M Tris, 0.1 M MgCl_2_, pH 9.0), 1 µL shrimp alkaline phosphatase (1000u/mL) and 100 µg lysate were mixed and filled up with H_2_O to a total volume of 100 µL. The reaction mixture was incubated overnight at 37 °C. As a positive control, the phosphorylated protein ovalbumin was used.

***Western blot analysis***

For each western blot analysis, 20 µg cell lysate was used. In order to denature proteins, 4 µL 5x Lämmli buffer (10 % (w/v) SDS, 50 % (v/v) glycerol, 25 % (v/v) β-mercaptoethanol, 0.5% (w/v) bromphenol blue, 315 mM Tris/HCl, pH 6.8) was added to 16 µL lysate solution and the samples were boiled 5 min at 95 °C. Afterwards, 20 µL of each sample was loaded onto a 7.5, 10 or 12.5 % SDS gel and proteins were separated according to their size by SDS-PAGE. Then, the separated proteins were transferred onto a membrane using a blotting sandwich moistened by blotting buffer (48 mM Tris, 39 mM glycine, 0.0375 % (m/v) SDS, 20 % (v/v) methanol), which was composed of one extra thick blot paper, the PVDF transfer membrane, the SDS-PAGE gel and again one extra thick blot paper. Before the protein transfer was carried out 45 min at 25 V using a Semi Dry Blotter (Bio-Rad), the transfer membrane was pre-incubated 5 min in methanol. In order to block non-specific binding sites, the membrane was incubated 60 min in blocking solution (0.5 g milk powder in 10 mL PBST (PBS + 0.5 % Tween)). Subsequently, 10 µL primary antibody with specificity for the protein of interest was added and the mixture was incubated 1 h at room temperature. The membrane was washed 3 times for 10 min with PBST before 1 µL of the secondary HRP antibody in 10 mL blocking solution was added. After 1 h of incubation at room temperature the membrane was washed again 3 times for 10 min with PBST. Then, 400 µL ECL Substrate and 400 µL peroxide solution were mixed and added to the membrane to stain the western blot. Finally, images of the western blot were taken by developing machine Amersham Imager 680 (GE Healthcare).

***Phos-tag gel***

In order to separate the unmodified from the post-translationally modified protein form, a Phos-tag gel was used. Compared to conventional SDS-gels, Phos-tag gels contain additionally Phos-tag^TM^ reagent and MnCl_2_. To pore a 1 mm thick 7.5 % Phos-tag separating gel, 1.25 mL 30 % acrylamide solution, 1.25 mL 1.5 M Tris pH 8.8 solution, 50 µL 5 mM Phos-tag^TM^ reagent, 50 µL 10 mM MnCl_2_, 50 µL 10 % SDS, 2.33 mL H_2_O, 5 µL TEMED and 25 µL 10 % APS were mixed. The stacking gel was prepared as for conventional SDS-gels by mixing 0.6 mL 30 % acrylamide solution, 1 mL 0.5 M Tris pH 6.8 solution, 40 µL 10 % SDS, 2.34 mL H_2_O, 4 µL TEMED and 20 µL 10 % APS. After the Phos-tag gel was polymerized, 20 µg of the lysates were loaded and dependent on the size of the protein of interest, the gel was run for 2 to 3 h at 150 V. Following the separation, the gel was washed 2 times 15 min with blotting buffer (48 mM Tris, 39 mM glycine, 0.0375 % (m/v) SDS, 20 % (v/v) methanol) supplemented with 10 mM EDTA to remove the manganese-ions. Subsequently, the gel was washed 15 min in blotting buffer before it was blotted as described in the western blot section.

***Calculation chemical proteomics***

MS Raw files were analysed using MaxQuant software 1.6.12.0 with the Andromeda search engine. Searches were performed against the Uniprot database for Homo sapiens (taxon identifier: 9606, March 2020). At least two unique peptides were required for protein identification. False discovery rate determination was carried out using a decoy database and thresholds were set to 1 % FDR both at peptide-spectrum match and at protein levels. LFQ quantification was used as described for each sample.

***Statistical analysis chemical proteomics***

Statistical analysis of the MaxQuant result table proteinGroups.txt was done with Perseus 1.6.10.43.

First, LFQ intensities were log_2_-transformed. Afterwards, potential contaminants as well as reverse peptides were removed. Then, the rows were divided into two groups - DMSO (control) and probe treated sample (sample). Subsequently, the groups were filtered for at least three valid values out of four rows in at least one group and the missing values were replaced from normal distribution. The -log_10_(p-values) were obtained by a two-sided one sample Student’s t-test over replicates with the initial significance level of p = 0.05 adjustment by the multiple testing correction method of Benjamini and Hochberg (FDR = 0.05) using the volcano plot function.

***Profile plots***

In order to visualize the changes of the LFQ-intensities during the iNGN differentiation, profile plots were used. For this, the mean (circle), the median (line) and the whiskers for outliers from *n* = 4 of each time point were prepared in Origin with box defined by 25^th^ and 75^th^ percentile.

***Data and Code availability***

Mass spectra from chemical proteomics and full proteomes are available via ProteomeXchange with identifier PXD023873. Reviewer account is accessible using the username: and password: AhqWCkko.
