## Supplemental Resource Table for "AMPylation is a specific lysosomal protein posttranslational modification in neuronal maturation"

**RESOURCES TABLE**

| REAGENT or RESOURCE | SOURCE | IDENTIFIER |
| --- | --- | --- |
| Antibodies | | |
| Goat polyclonal anti-mouse IgG, AF488-linked | Thermo Fisher Scientific | Cat# A-11001; RRID:AB_2534069 |
| Goat polyclonal anti-rabbit IgG, AF488-linked | Thermo Fisher Scientific | Cat# A-11008; RRID: AB_143165 |
| Goat polyclonal anti-rabbit IgG, HRP-linked | Sigma-Aldrich | Cat# A6667; RRID: AB_258307 |
| Rabbit polyclonal anti-ABHD6 | Thermo Fisher Scientific | Cat# PA5-38999; RRID AB_2555591 |
| Rabbit polyclonal anti-ACP2 | Thermo Fisher Scientific | Cat# PA5-29961; RRID: AB_2547435 |
| Rabbit polyclonal anti-COX IV | Thermo Fisher Scientific | Cat# ab16056; RRID: AB_443304 |
| Rabbit polyclonal anti-HSPA5 | Thermo Fisher Scientific | Cat# PA5-34941;  RRID: AB_2552290 |
| Mouse monoclonal anti-KDEL (10C3) | Millipore | Cat# 420400;  RRID: AB_212090 |
| Mouse monoclonal anti-LAMP2 (H4B4) | DSHB | Cat# N/A; RRID:  [AB_528129](http://antibodyregistry.org/AB_528129) |
| Rabbit polyclonal anti-PLD3 | Sigma-Aldrich | Cat# HPA012800; RRID: AB_1855330 |
| Mouse monoclonal anti-TUBB3 | Sigma-Aldrich | Cat# T8660  RRID: AB_477590 |
| Guinea pig polyclonal anti-DCX | Millipore | Cat# AB2253  RRID: AB_1586992 |
| Rabbit polyclonal anti-TBR1 | Millipore | Cat# AB31940  RRID: AB_2200219 |
| Mouse polyclonal anti-MAP2 | Millipore | Cat# AB5392  RRID: AB_2138153 |
| Mouse monoclonal anti-Neun | Millipore | Cat# MAB377  RRID: AB_2298772 |
| Goat polyclonal anti-mouse IgG2b, AF647-linked | Thermo Fisher Scientific | Cat# A-21242  RRID: AB_2535811 |
| Goat polyclonal anti-rabbit IgG, AF546-linked | Thermo Fisher Scientific | Cat# A-11010  RRID: AB_2534077 |
| Goat polyclonal anti-guinea pig IgG, AF647-linked | Thermo Fisher Scientific | Cat# A-21450  RRID: AB_2735091 |
| Goat polyclonal anti-mouse IgG, AF546-linked | Thermo Fisher Scientific | Cat# A-21123  RRID: AB_2535765 |
| Mouse monoclonal anti-S100beta | Sigma-Aldrich | Cat# S2532  RRID: AB_477499 |
| Rabbit polyclonal anti-FGFR3 | Santa Cruz | Cat# sc-123  RRID: AB_631511 |
| Goat anti-mouse IgG1, Alexa Fluor 488 | Thermo Fisher Scientific | Cat# A-21121  RRID: AB_2535764 |
| Bacterial and Virus Strains | | |
| Biological Samples |  |  |
| Chemicals, Peptides, and Recombinant Proteins | | |
| Acetone (HPLC grade) | VWR chemicals | Cat# 200067.320; CAS: 67-64-1 |
| Acetonitrile (LC-MS grade) | Fisher Scientific | Cat# A955-212; CAS: 75-05-8 |
| **Alanyl-Glutamine** | Sigma-Aldrich | Cat# G8541, CAS: 39537-23-0 |
| Ammoniumperoxodisulfat (APS) | Sigma-Aldrich | Cat# 09913; CAS: 7727-54-0 |
| β-Mercaptoethanol | Sigma-Aldrich | Cat# M3148; CAS: 60-24-2 |
| Biotin-PEG_3_-N_3_ | Carbosynth | Cat# FA34890 CAS: 875770-34-6 |
| Bromphenol blue | Fluka | Cat# 32712 CAS: 115-39-9 |
| BSA | AppliChem | Cat# A6588; CAS: 9048-46-8 |
| Coomassie Blue R-250 | Fluka | Cat# 27816; CAS: 6104-59-2 |
| CuSO_4_ x 5 H_2_O | Acros | Cat# 10627162, 7758-99-8 |
| ddH2O (LC-MS grade) | Honeywell | Cat# 15665350 CAS: 732-18-5 |
| DMSO | Sigma-Aldrich | Cat# D4540, CAS: 67-68-5 |
| Dithiothreitol | VWR (AppliChem) | Cat# A2948, CAS: 3483-12-3 |
| Formic Acid (LC-MS grade) | Fisher Scientific | Cat# A117; CAS: 64-18-6 |
| Hepes | Carl Roth | Cat# HN77.5; CAS: 7365-45-9 |
| Iodacetamide | Sigma-Aldrich | Cat# I6125; CAS: 144-48-9 |
| KCl | AppliChem | Cat# A2939; CAS: 7447-40-7 |
| KH_2_PO_4_ | Sigma-Aldrich | Cat# P9791; CAS: 7778-77-0 |
| Na_2_HPO_4_ | Sigma-Aldrich | Cat# T876; CAS: 7558-79-4 |
| Na_2_SeO_3_ | Sigma-Aldrich | Cat# S5261; CAS: 10102-18-8 |
| NaCl | Bernd Kraft GmbH | Cat# KRAF04160; CAS: 7647-14-5 |
| NP40 | Sigma-Aldrich | Cat# 74385; CAS: 9016-45-9 |
| Methanol (LC-MS grade) | Fisher Scientific | Cat# A456; CAS: 67-56-1 |
| Powdered milk | AppliChem | Cat# A0830; CAS: 999999-99-4 |
| SDS | AppliChem | Cat# A2572; CAS: 151-21-3 |
| Sodium Deoxycholate | Sigma-Aldrich | Cat# 30970; CAS:  302-95-4 |
| TAMRA-N_3_ | Baseclick | CAT# BCFA-008-1 |
| TBTA | TCI | Cat# T2993; CAS: 510758-28-8 |
| TCEP | Carbosynth | Cat# FT01756; CAS: 51805-45-9 |
| TEAB (1 M) | Sigma-Aldrich | Cat# T7408, CAS: [15715-58-9](https://www.sigmaaldrich.com/catalog/search?term=15715-58-9&interface=CAS%20No.&N=0&mode=partialmax&lang=de&region=DE&focus=product) |
| TEMED | Sigma-Aldrich | Cat# T9281;CAS: 110-18-9 |
| Thiourea | Merck | Cat# 107979; CAS: 62-56-6 |
| Tris-base | Fisher Scientific | Cat# 10724344; CAS: 77-86-1 |
| Trypan Blue | Fisher Scientific | Cat# 11538886 |
| Tween^®^ 20 | VWR (AppliChem) | Cat# A4974; CAS: 9005-64-5 |
| Urea | AppliChem | Cat# A1049, CAS: 57-13-6 |
| DPBS (1x) | Sigma-Aldrich | Cat# D8357 |
| DMEM (1x) | Sigma-Aldrich | Cat# D6546 |
| Ham's F-12 w/o L-Glu | Sigma-Aldrich | Cat# N4888 |
| cOmplete® Protease Inhibitor | Sigma-Aldrich | Cat# 05056489001 |
| Normal Goat Serum | Biozol | Cat# VEC-S-1000 |
| FCS | Thermo Fisher Scientific | Cat# [A3840001](https://www.thermofisher.com/order/catalog/product/A3840001) |
| hHolo-Transferrin | Sigma-Aldrich | Cat# 616424; CAS: 11096-37-0 |
| hFGF-2 | MACS Miltenyi Biotec | Cat# 130-104-921 |
| hInsulin | BioXtra | Cat# I9278, CAS: 11061-68-0 |
| hTGF- β1 | PeproTech |  |
| Geltrex | Thermo Fisher Scientific | Cat# A1413201 |
| Immobilon® Western HRP Sustrate | Merck Millipore | Cat# WBKLS0500 |
| Color Prestained Protein Standard, Broad Range (10-250 kDa) | New England BioLabs | Cat# P7719S |
| Pen-Strep | Sigma-Aldrich | Cat# P0781 |
| Rotiphorese.Gel 30 (37, 5:1) | Carl Roth | Cat# 3029.1 |
| TrypLE Express | Thermo Fisher Scientific | Cat# 12604013 |
| Trypsin | Promega | Cat# V5113 |
| Thiazovivin | Merck Millipore | Cat# 420220; CAS: 1226056-71-8 |
| NeuroBrew-21 | Miltenyi Biotech | Cat# 130-093-566 |
| 4,6‐diamidino‐2‐phenylindole | Sigma-Aldrich | Cat# D9542 |
| Accutase | Stem Cell Technologies | Cat# 07920 |
| Ascorbic acid | Sigma-Aldrich | Cat# A92902; CAS: [50-81-7](https://www.sigmaaldrich.com/catalog/search?term=50-81-7&interface=CAS%20No.&N=0&mode=partialmax&lang=de&region=DE&focus=product) |
| B27-supplement (minus vitamin A) | Thermo Fisher Scientific | Cat# 12587010 |
| BDNF | Peprotech | Cat# 450-02 |
| dcAMP | Sigma-Aldrich | Cat# D1256; CAS: [93839-95-3](https://www.sigmaaldrich.com/catalog/search?term=93839-95-3&interface=CAS%20No.&N=0&mode=partialmax&lang=de&region=DE&focus=product) |
| GDNF | Peprotech | Cat# 450-10 |
| Laminin | Sigma-Aldrich | Cat# L2020; CAS: 114956-81-9 |
| Matrigel® Basement Membrane Matrix, LDEV-free | Corning® | Cat# 354234 |
| Minimum essential medium/non-essential amino acid | Thermo Fisher Scientific | Cat# 11140050 |
| mTESR1 medium | Stem Cell Technologies | Cat# 85850 |
| N2-supplement | Thermo Fisher Scientific | Cat# 17502048 |
| Neurobasal medium | Thermo Fisher Scientific | Cat# 21103049 |
| Polyornithine | Sigma-Aldrich | Cat# P4957 |
| Rock inhibitor Y-27632(2HCl) | Stem Cell Technologies | Cat# 72304 |
| Astrocyte Media | ScienCell | Cat# 1801 |
| Recombinant Human LIF | Alomone Labs | Cat# L-200 |
| Recombinant Human FGF-basic | Peprotech | Cat# AF-100-18B |
| Recombinant Human EGF | Peprotech | Cat# AF-100-15 |
| Recombinant Human Noggin | Peprotech | Cat# 120-10C |
| Recombinant Human PDGF-AA | R&D Systems | Cat# 221-AA |
| Accutase | Thermo Fisher Scientific | Cat# A1110501 |
| Collagenase | Stem Cell Technologies | Cat# 07909 |
| Critical Commercial Assays | | |
| Pierce® BCA Protein Assay Kit | Thermo Fisher Scientific | Cat# 23225 |
| Deposited Data | | |
| MS raw data and calculation results | ProteomeXchange | PXD023873 |
| Experimental Models: Cell Lines | | |
| Human: HeLa |  | RRID: CVCL_0030 |
| Human: SH-SY5Y |  | RRID: CVCL_0019 |
| Human: iNGNs | Volker Busskamp, CRTD Dresden |  |
| Human: iPSCs | Dr.Micha Drukker  HHZ Munich | HMGU No 1  RRID: CVCL_YT30 |
| Experimental Models: Organisms/Strains | | |
| Oligonucleotides | | |
| Recombinant DNA | | |
| Software and Algorithms | | |
| MaxQuant | Cox et al., 2014. | https://www.maxquant.org/download_asset/maxquant/latest |
| Perseus | Tyanova et al., 2016. | https://maxquant.net/download_asset/perseus/latest |
| Origin | N/A | https://www.originlab.com/ |
| Other | | |
